# Application of Attention and Graph Transformer-Based Approaches for RNA Biomarker Discovery in Metabolically-Associated Fatty Liver Disease (MAFL/NASH)

**DOI:** 10.1101/2023.11.05.565710

**Authors:** Aashish Cheruvu, Daniel Zezulinski, Aejaz Sayeed

## Abstract

The prevalence of nonalcoholic fatty liver disease (NAFLD) and nonalcoholic steatohepatitis (NASH) in the United States has reached epidemic proportions, increasing the risk of liver cirrhosis and cancer. Current methods of diagnosis for NAFLD/NASH are invasive and costly, motivating the need for genetic “RNA” biomarkers detectable in a blood sample. In this study, explainable artificial intelligence (XAI) techniques are employed to increase the interpretability of the deep learning models in detecting the potential mRNA biomarker candidates for NAFLD/NASH. Nine RNA datasets (∼1000 patients) with NAFLD/NASH were collected from the Gene Expression Omnibus. After conducting a differential gene expression analysis to reduce the dimensionality of the expression data, single-head and multi-head attention models were compared to baseline machine learning models in their ability to classify patients as NAFLD/NASH/healthy. XAI methods, including L1 regularization on baseline models and analysis of the internal attention matrix of the attention models, were utilized to identify biomarker candidates based on the relative importance of genes. The attention models achieved superior performance (accuracy: 67.5%) compared to the baseline models (Negative Binomial Linear Discriminant Analysis-62.64%; Poisson Linear Discriminant Analysis with Power Transformation – 58.24%). The top 17 and top 20 XAI-identified biomarkers with the baseline machine learning algorithms and the attention-based models respectively were then evaluated in lab. Preliminary data from in-lab validation confirmed upregulation of MT-ND3, HLA-B, APOC-1, and APOL-1 in NAFLD/NASH patients. Attention models have shown promise in identifying expression-based mRNA biomarkers and accurately diagnosing patients with NAFLD/NASH.

## 1. INTRODUCTION

Early detection of cancer can have a significant impact on patient outcomes, as it can increase the chances of successful treatment and improve survival rates. Cancer that is detected at an early stage is often more treatable and more likely to respond to treatment. Conversely, cancer detected at a later stage may have already spread or advanced, making it more challenging to treat and lowering the chances of successful outcomes. Hepatocellular carcinoma (HCC), the most common type of liver cancer, is particularly challenging to detect early, as it is often asymptomatic in its early stages^1^. This means that patients may not experience any symptoms until the cancer has progressed to a more advanced stage. Consequently, early detection is crucial in improving patient outcomes.

Nonalcoholic associated fatty liver disease (NAFLD), also known as metabolically associated fatty liver disease (MAFLD), is an early-stage liver disease that is increasingly prevalent in the United States^2^. It affects an estimated 24% of U.S. adults, and early detection is important in preventing its progression to nonalcoholic steatohepatitis (NASH), a more severe form of the disease that can lead to inflammation, liver cell damage, and scarring/fibrosis of the liver. Over time, this scarring can result in the development of liver cirrhosis, liver failure, and even hepatocellular carcinoma (HCC), a type of liver cancer. However, diagnosing NAFLD/NASH is challenging, as few symptoms are visible in the early stages, and the current gold standard for diagnosis, a liver biopsy, is expensive, invasive, and carries a risk of serious complications. Therefore, identifying new biomarkers and developing non-invasive diagnostic tools that can differentiate between healthy individuals and those with NAFLD/NASH could lead to earlier detection of the disease and better patient outcomes.

Traditionally, protein biomarkers have been used in the diagnosis of diseases including cancer, heart disease, and infectious diseases. These biomarkers are proteins that are produced by cells in response to a disease or condition and can be detected in blood, urine, or tissue samples. However, there are limitations to the use of protein biomarkers in disease diagnosis. For example, some protein biomarkers may not be specific to a particular disease or condition and may also be produced in response to other factors such as inflammation or injury. Additionally, some protein biomarkers may be present at very low levels, making them difficult to detect, or may vary widely between individuals, making it difficult to establish a reliable cutoff for disease diagnosis. As a result, there is growing interest in the use of other biomarkers, such as RNA biomarkers, which may offer improved sensitivity and specificity in disease diagnosis^3^. RNA biomarkers are molecules that are produced by cells in response to disease or other conditions and can be detected in blood or other body fluids. These biomarkers may provide more accurate and reliable information about disease status and progression, leading to earlier diagnosis and more effective treatment.

RNA datasets are large and complex, typically containing tens of thousands of features (genes) that need to be analyzed in relation to a smaller number of samples, which can be in the hundreds^4^. This creates a statistical problem known as the “p >> n” problem, where the number of features is much greater than the number of samples. This makes it difficult to analyze the data using traditional statistical methods, which may be prone to overfitting or underfitting the data and may not be able to effectively identify patterns or associations between features and samples. To overcome these challenges, machine learning techniques have been developed to analyze RNA data. Machine learning is a type of artificial intelligence that involves developing algorithms that can learn from data and make predictions or decisions based on that data. These algorithms can be trained to identify patterns and relationships in large and complex datasets. The use of machine learning techniques for the analysis of RNA data has several advantages, including their ability to identify patterns and relationships that may not be apparent using traditional statistical methods, their ability to handle large and complex datasets, and their ability to learn from new data and adapt to changing conditions.

Explainable Artificial Intelligence (XAI) is an emerging field that focuses on developing machine learning models that are both accurate and explainable^5^. In many applications, machine learning models need to be both accurate and interpretable, particularly in fields such as healthcare and finance, where decisions can have significant consequences. There is often a trade-off between performance and explainability in machine learning models. Models that prioritize performance may be complex and difficult to understand, while models that prioritize explainability may sacrifice some accuracy. Therefore, XAI seeks to strike a balance between these two factors, by developing models that are both accurate and interpretable. Explainability is particularly important in fields such as healthcare, where decisions made by machine learning models can have significant consequences for patients. For example, a model that predicts the likelihood of a patient having a particular disease needs to be both accurate and interpretable so that clinicians can understand the basis for the model’s prediction and make informed decisions about patient care.

Attention mechanisms are an important component of XAI^6^. In recent years, attention-based models have become an essential tool in healthcare, enabling significant progress in medical research and improving patient outcomes. Attention is a mechanism that allows machine learning models to selectively focus on specific parts of the data while ignoring irrelevant information. This mechanism has proven to be highly effective in improving the accuracy of predictions, making it a key component of state-of-the-art machine learning models such as transformers. Transformers are a type of neural network that can process sequential data with exceptional accuracy, making them well-suited to many healthcare applications. These models use self-attention mechanisms to selectively focus on the most relevant parts of the input data, allowing them to learn complex patterns and relationships in the data. Some notable examples of transformers in healthcare include ChatGPT and Stable Diffusion.

The objective of this study is to identify new biomarker candidates for Nonalcoholic Associated Fatty Liver Disease (NAFLD)/Nonalcoholic Steatohepatitis (NASH) using publicly available Gene Expression Omnibus (GEO) datasets of NAFLD/NASH patients and healthy controls. The study employs both traditional machine learning-based algorithms and advanced deep learning techniques, leveraging state-of-the-art explainable AI (XAI) methodologies such as attention mechanisms. These advanced methods aim to uncover previously unrecognized patterns and relationships within the large and complex datasets, leading to the discovery of novel biomarkers for early detection and diagnosis of NAFLD/NASH.

## 2. RELATED WORK

In the realm of cancer diagnosis and prognosis, a systematic review has shed light on the state-of-the-art computational methods, including machine learning and deep learning, which are pivotal in the discovery of biomarkers from complex multi-omics data^15^. This underscores the importance of advanced feature selection strategies and the availability of sophisticated tools that are shaping the current landscape of biomarker research. Moreover, the application of machine learning extends to human microbiome studies, where its utility in diagnostics, prognostics, and therapeutic contexts has been explored, highlighting the algorithmic flexibility to adapt to various biological data types^16^. In the field of machine vision, attention mechanisms have been introduced and discussed, providing insights into how these could be applied to biomarker identification, particularly through image-based data analysis^17^. Studies have also demonstrated the effectiveness of machine learning in identifying novel predictive biomarkers for lung diseases, suggesting a broader potential for ML in uncovering biomarkers across a spectrum of diseases^18^. In medical imaging, attention-based convolutional neural networks have been proposed for brain tumor segmentation, indicating the potential of attention mechanisms to enhance the interpretability of complex data, a concept that can be translated into the identification of RNA biomarkers for diseases like NAFLD/NASH^19^. Furthermore, traditional feature selection techniques have been employed for cancer classification and biomarker discovery, while research on human muscle transcriptomic data has utilized machine learning to confirm established mechanisms of aging and identify new biomarkers^20,21^. The concept of dual cross-attention learning has been proposed to reduce misleading attentions and discover more complementary parts for recognition, an approach that could inspire similar methodologies in biomarker identification^22^. The pioneering application of machine learning for ADHD detection using pupillometric biomarkers and the identification of dysregulated miRNAs as reliable biomarkers for gastric cancer using machine learning highlight the transformative power of these computational techniques in the field of medical diagnostics^23^. Collectively, these studies not only demonstrate the growing interest but also the tangible potential in employing machine learning and attention mechanisms for the identification and validation of biomarkers, offering a promising avenue to enhance the accuracy and interpretability of diagnostic models in various diseases, including NAFLD/NASH^24^.

## 3. MATERIALS AND METHODS

### 2.1. *In Silico* analysis

#### 2.1.1 Computational methods

In this study, Python and R were the primary programming languages used for data preparation, analysis, and visualization. The Google Cloud platform was used for cloud computing, which provided access to high-performance computing resources, such as CPUs, GPUs, and TPUs, for model building and evaluation. To facilitate data visualization, the matplotlib library was used, while the ‘pandas’ library was used for data frame object and data manipulation. The ‘NumPy’ library was utilized for random sampling and fast mathematical operations, and ‘scikit-learn’ was used to calculate the metrics required for this project. In addition, ‘PyTorch’, a deep learning library with a dynamic graph structure, automatic gradient calculation, and GPU acceleration, was used to develop and evaluate the deep learning models. The combination of these tools and libraries helped to optimize the performance of the machine learning models, ultimately leading to improved accuracy and better insights into the complex data.

#### 2.1.2 Dataset Acquisition

Our study aims to explore the transcriptome profiles of liver biopsy datasets obtained from patients diagnosed with non-alcoholic steatohepatitis (NASH) or non-alcoholic fatty liver disease (NAFLD). To accomplish this, we retrieved data from the National Center for Biotechnology Information’s (NCBI) Gene Expression Omnibus (GEO), a comprehensive database of publicly available gene expression data^7^. Identified 9 datasets that met inclusion criteria, which included a total of 737 samples from patients across 7 countries (Table 1). These 9 datasets represent a diverse range of biological, clinical, and technical heterogeneity, providing a comprehensive snapshot of the disease across different populations. They include samples from both adolescents and adults, with or without comorbidities, and were profiled using a variety of commercial microarrays and RNA sequencing platforms. One strength of these datasets is the inclusion of healthy controls, which were representative of real-world heterogeneity. They included individuals with normal weight, healthy obese individuals, and those suspected of having NAFLD. This allows for a more accurate comparison between the diseased and healthy states and provides a better understanding of the disease pathology. Overall, the use of these 9 diverse datasets provides a robust and comprehensive platform for the investigation of NASH and NAFLD transcriptome profiles, enhancing our understanding of the disease and potential therapeutic targets.

**Table 1.** Liver biopsy datasets collected from GEO Database. Datasets included in analysis include data across multiple countries, ages, and technical variation in gene expression platforms. In the NASH/NAFLD and Healthy Control (HC) columns. N indicates the number of samples in the relevant group.

| **Dataset ID** | **Country** | **Age** | **NAFLD/NASH** | **HC** | **Available phenotypes** | **Platform** |
| --- | --- | --- | --- | --- | --- | --- |
| E-MEXP-3291 | US | 16–70 | 16/10 | 19 | Sex, age | GPL6244 Affymetrix Human Gene 1.0 ST Array |
| GSE61260 | Germany | 20–86 | 24/23 | 62 | Sex, age, BMI | GPL11532 Affymetrix Human Gene 1.1 ST Array |
| GSE66676 | US | 13–20 | 7/26 | 34 | Sex, age, BMI, histology, HDL, LDL, cholesterol, triglycerides | GPL6244 Affymetrix Human Gene 1.0 ST Array |
| GSE126848 | Denmark | 18-70 | 16/15 | 26 | Sex | GPL18573 RNAseq, Illumina NextSeq 500 |
| GSE135251 | France, Germany, Italy and the UK | 54 ± 11.87 | 53/153 | 10 | Sex, age, BMI, histology, HDL, LDL, cholesterol, triglycerides | Illumina NextSeq 500 |
| GSE207310 | Denmark | 18-70 | 10/15 | 5 | Sex, age, BMI, histology, HDL, LDL, cholesterol, triglycerides | Illumina NovaSeq6000 platform |
| GSE192959 | Global | 67-75 | 48 | 0 | Sex, age, BMI | Illumina NextSeq 500 |
| GSE193066 | Global | 42-62 | 106 | 0 | Sex, age, BMI | Illumina NextSeq 500 |
| GSE193080 | Global | 65-77 | 59 | 0 | Sex, age, BMI | Illumina NextSeq 500 |

#### 2.1.3 Data Preprocessing

This study analyzed multiple datasets to discover RNA biomarkers, which involved the normalization of data from different sources to remove site variation. The RNA-seq data (from GSE135251, GSE126848, GSE207310) was normalized using Transcripts Per Million (TPM) normalization across genes, while the microarray data (from GSE66676, GSE61260, E-MEXP-3291) was normalized using Robust Multi-Array Average (RMA) across genes to remove site variation. In addition, the Relative Log Expression (RLE) normalized count data from GSE192959, GSE193066, and GSE193080 was back transformed into raw count data and then normalized using TPM. By normalizing the data in this way, reduced the effects of variability due to the different sources and provide more accurate and reliable results. 12

#### 2.1.4 Model architecture

##### 2.1.4.1 Baseline Machine Learning methods

A total of 737 mRNA-Seq datasets, constituted the compiled machine learning (ML) dataset for further classification and feature selection. Raw gene counts generated for each dataset were processed and analyzed in R v4.0.2 with the Bioconductor package MLSeq v2.6.0 https://github.com/dncR/MLSeq)^8^. The objectives of the ensuing ML analysis were to develop ML models that would (1) accurately “classify” an mRNA-Seq dataset within the 9 datasets and (2) extract genes and gene sets or “features” that accurately assign an mRNA-Seq dataset to its disease (NAFLD/NASH) or healthy class.

The classification and biomarker identification method in brief can be summarized as, offset values of one were added to the count matrix to reduce the likelihood of convergence in model fitting and to reduce bulk sparsity. Genes with a minimum count-per-million of 0.5 in three or more mRNA-Seq libraries were retained for analysis. Library normalization was performed with the DESeq median ratio approach, using default settings. The resulting ML dataset was stratified into a training and testing set (70% and 30%, respectively), using controls as the comparative baseline (i.e., class statement).

Model validation and parameter optimization were evaluated using fivefold, 10 repeats with non-exhaustive cross validation. Six ML models were utilized for classification and/or significant gene selection: sparse Poisson linear discriminant analysis, with and without a power transformation (PLDA, PLDA2), negative binomial linear discriminant analysis (NBLDA), sparse voom-based nearest shrunken centroids (VNSC), support vector machine (SVM) (https://cran.r-project.org/web/packages/caret/caret.pdf). Models were evaluated with confusion matrices and performance metrics provided by the MLSeq package. Feature selection from sparse classifier models was set to a maximum of 2000 genes, based on maximum variance filtering. Sparse classifier models (PLDA, PLDA2, VNSC, and PAM), which generate lists of a select number of significant genes used for model decision and classification, were manually designated as the top models for each test set based on highest associated balanced accuracy and Kappa statistic; if two or more models were equal, gene lists would be merged. Performance metric calculations are defined by Goksuluk and colleagues. Balanced accuracy, the combined average of sensitivity and specificity, was a prioritized metric due to imbalance between challenged and control cattle and potential for skewed results when evaluating sensitivity and specificity alone. Further information regarding workflow parameters, model building, and optimization are found in the package vignette and associated GitHub repository mirror (https://bioconductor.org/packages/release/bioc/html/MLSeq.html; https://github.com/dncR/MLSeq).

Multidimensional scaling was applied to the gene count matrix with plotMDS, using pairwise distances of the top 500 genes based on variance68. Heatmaps of the unique gene classifiers identified across etio ogic test sets were generated with the Bioconductor package pheatmap v1.0.1269, utilizing Ward’s method of unsupervised hierarchical clustering on Euclidean distances and Pearson correlation coefficients for samples and genes, respectively. Color scaling for all packages was performed with the Bioconductor package viridis v0.5.1 70 to allow ease of visual interpretation for individuals with color blindness.

##### 2.1.4.2 Explainable Artificial Intelligence (XAI) -Single-head Attention/Multi-head attention methods

Attention and multi-head attention are both important mechanisms in machine learning that allow models to selectively focus on relevant parts of the input data^9^. While both mechanisms serve similar purposes, they differ in how they process and weight the input data.

##### 2.1.4.3 Attention architecture

The attention mechanism works by transforming the input data into Query (Q), Key (K), and Value (V) matrices^10^. The Q matrix represents the information that the model wants to retrieve from the input, while the K matrix represents the input itself. The model computes attention weights between Q and K, which are used to weight the values in the V matrix that are used to generate the output. The attention operator can be illustrated as shown in Figure 3.

**Figure 2.**
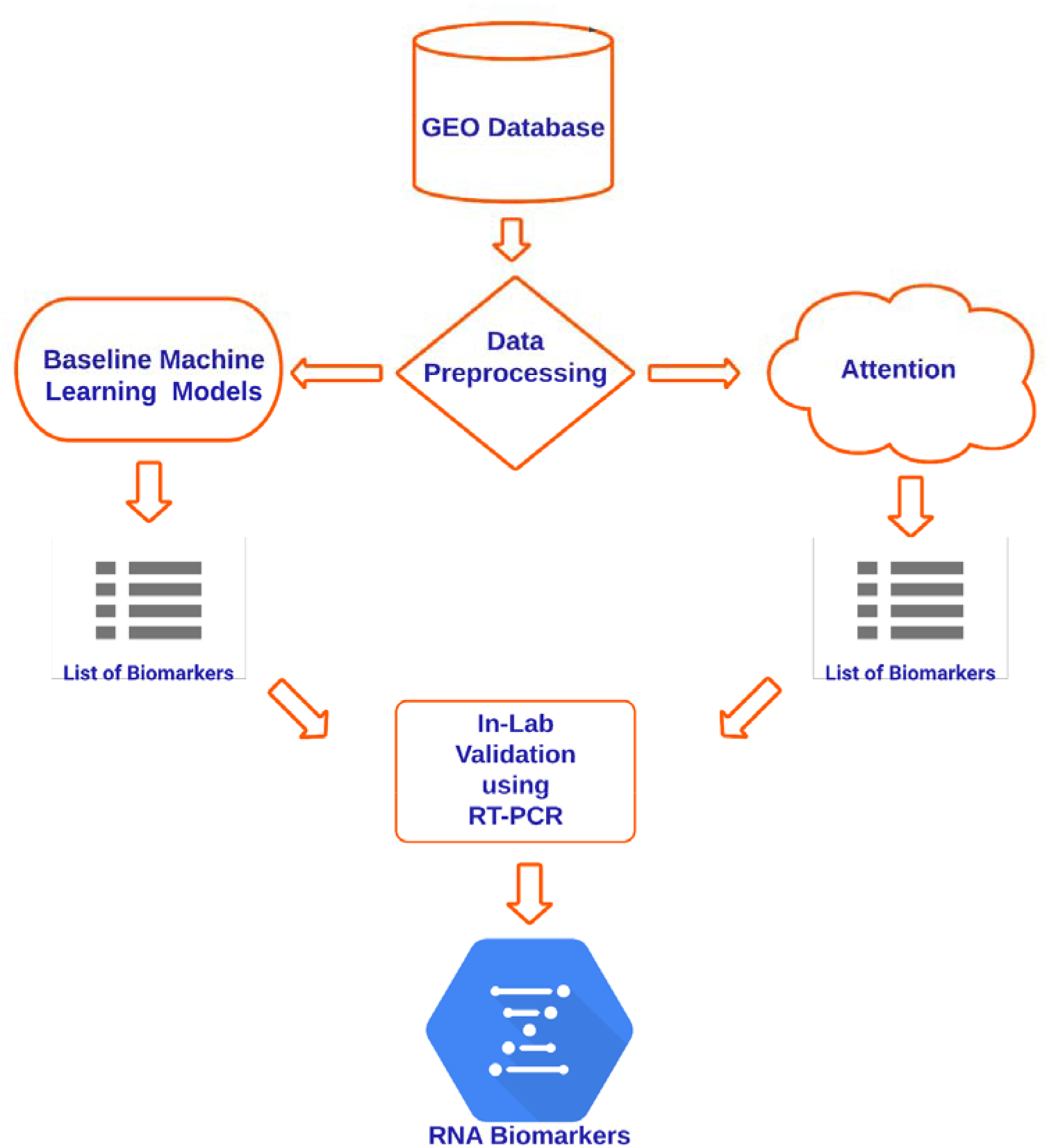
The following flow chart was used in finding and validating biomarkers.

**Figure 3:**
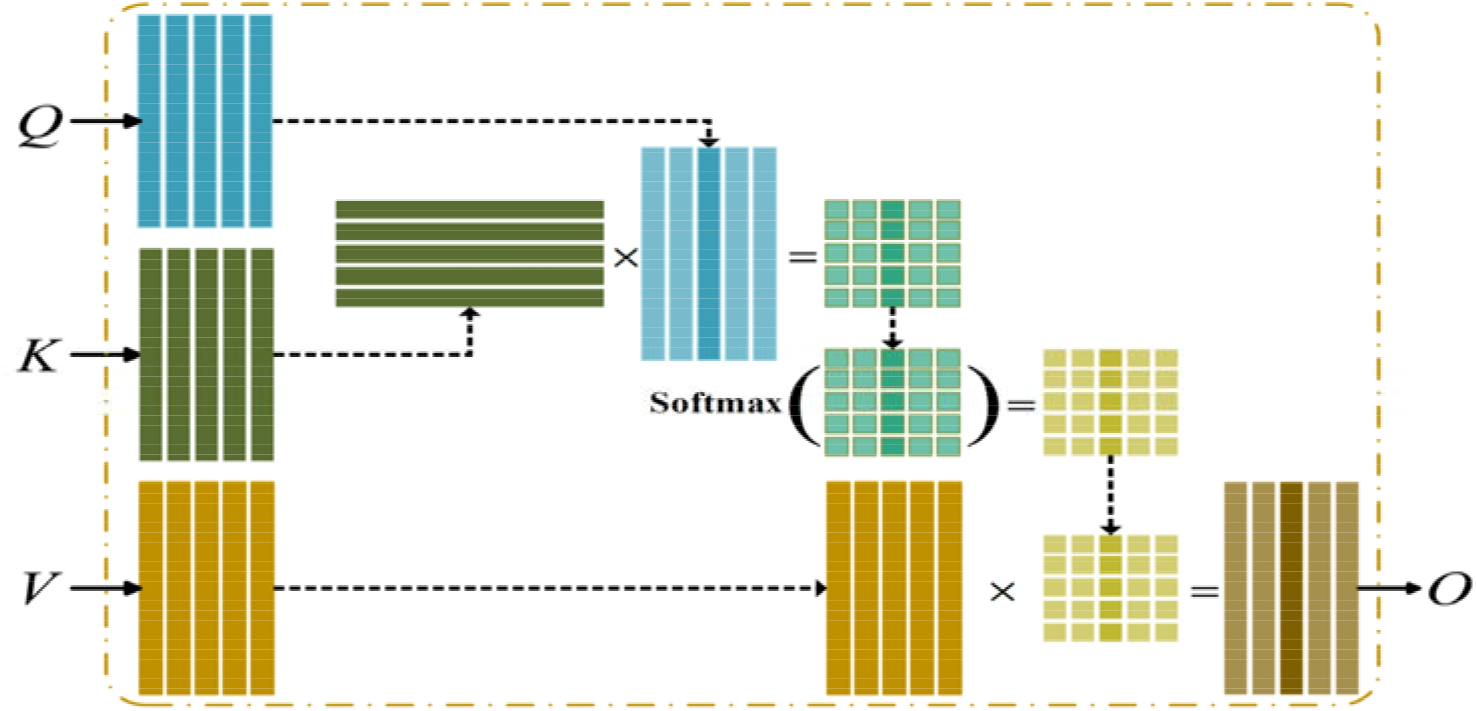
Illustration of attention operator.

x denotes matrix multiplication, and Softmax (•) is the column-wise softmax operator. Q, K, V are input matrices. A similarity score is computed between each query vector as a column of Q and each key vector as a column in K. Softmax (•) normalizes these scores and makes them sum to 1. Multiplication between normalized scores and the matrix V yields the corresponding output vector.

Attention mechanism can be represented mathematically as

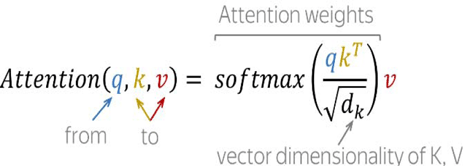

The attention mechanism is useful in applications where the input data contains complex relationships or dependencies between different parts of the input. By selectively focusing on relevant parts of the input, the attention mechanism can significantly improve the accuracy and interpretability of machine learning models.

##### 2.1.4.4 Multi-head Attention architecture

Multi-head attention is an extension of the attention mechanism that allows for parallel processing of Q, K, and V^11^. Instead of computing a single attention score for each Q-K pair, the multi-head attention mechanism computes multiple attention scores, or “heads,” in parallel. Each head of the multi-head attention mechanism has its own set of Q, K, and V weight matrices, which are learned during the training process.

The multi-head attention operator can be illustrated as shown in Figure 4.

**Figure 4:**
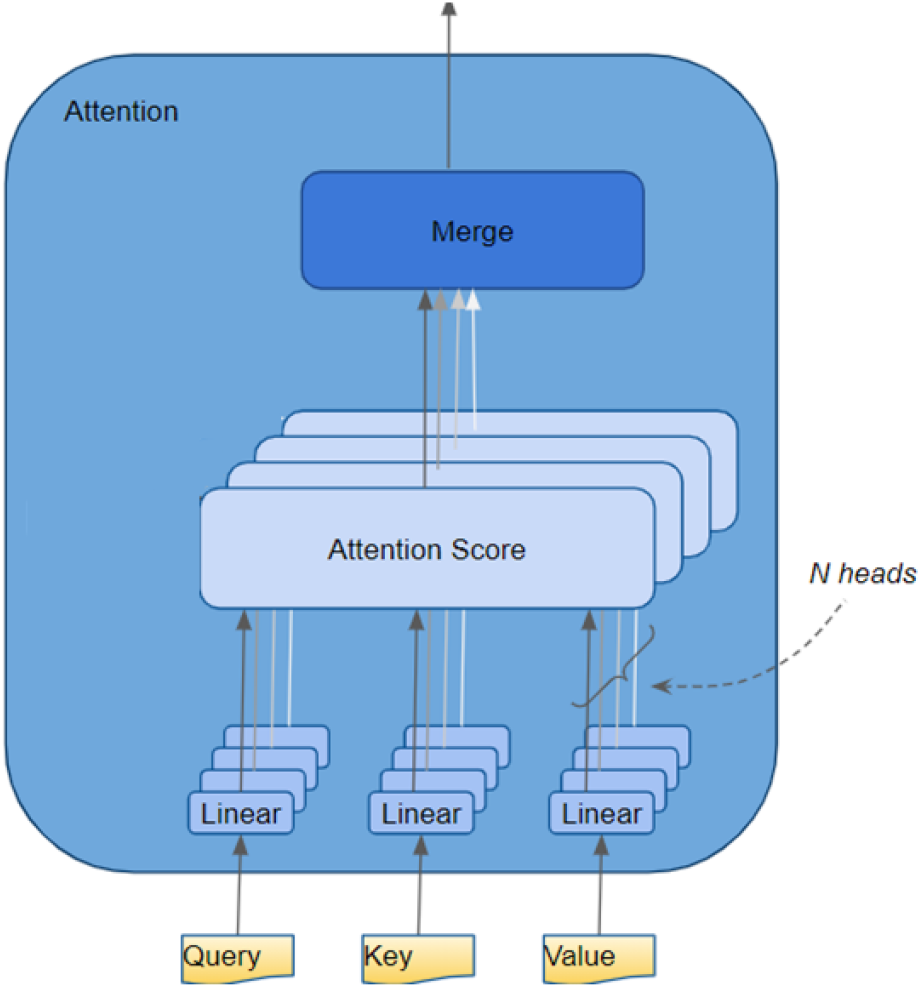
Multi-head Attention architecture.

The use of multi-head attention allows the model to capture more complex relationships and interactions within the input data, as each head can attend to different aspects of the input. This can lead to improved performance on tasks such as language modeling and machine translation, where capturing complex relationships between different parts of the input is crucial.

Overall, both attention and multi-head attention are important mechanisms in machine learning that allow models to selectively focus on relevant parts of the input data. While attention is useful in many applications, multi-head attention can capture more complex relationships and interactions within the input data, leading to improved performance on certain tasks. By allowing models to focus on relevant parts of the input data, attention and multi-head attention can significantly improve the accuracy and interpretability of machine learning models.

##### 2.1.4.5 Graph Attention Networks

Graph data refers to any data that can be represented in the form of a graph structure, where nodes represent entities and edges represent relationships between them. Examples of graph data include social networks, biological networks, traffic networks, and citation networks. Graph data is different from traditional data such as tabular data, text data, and image data, as it has no inherent order or sequence and requires specialized techniques to analyze.

Graph Neural Networks (GNNs) are a class of neural networks that are designed to process graph data^12^. GNNs learn node representations by aggregating information from neighboring nodes and using this information to update the representation of the current node. GNNs have shown promising results in various graph-related tasks such as node classification, link prediction, and graph classification. The architecture of GAT can be represented as shown below:

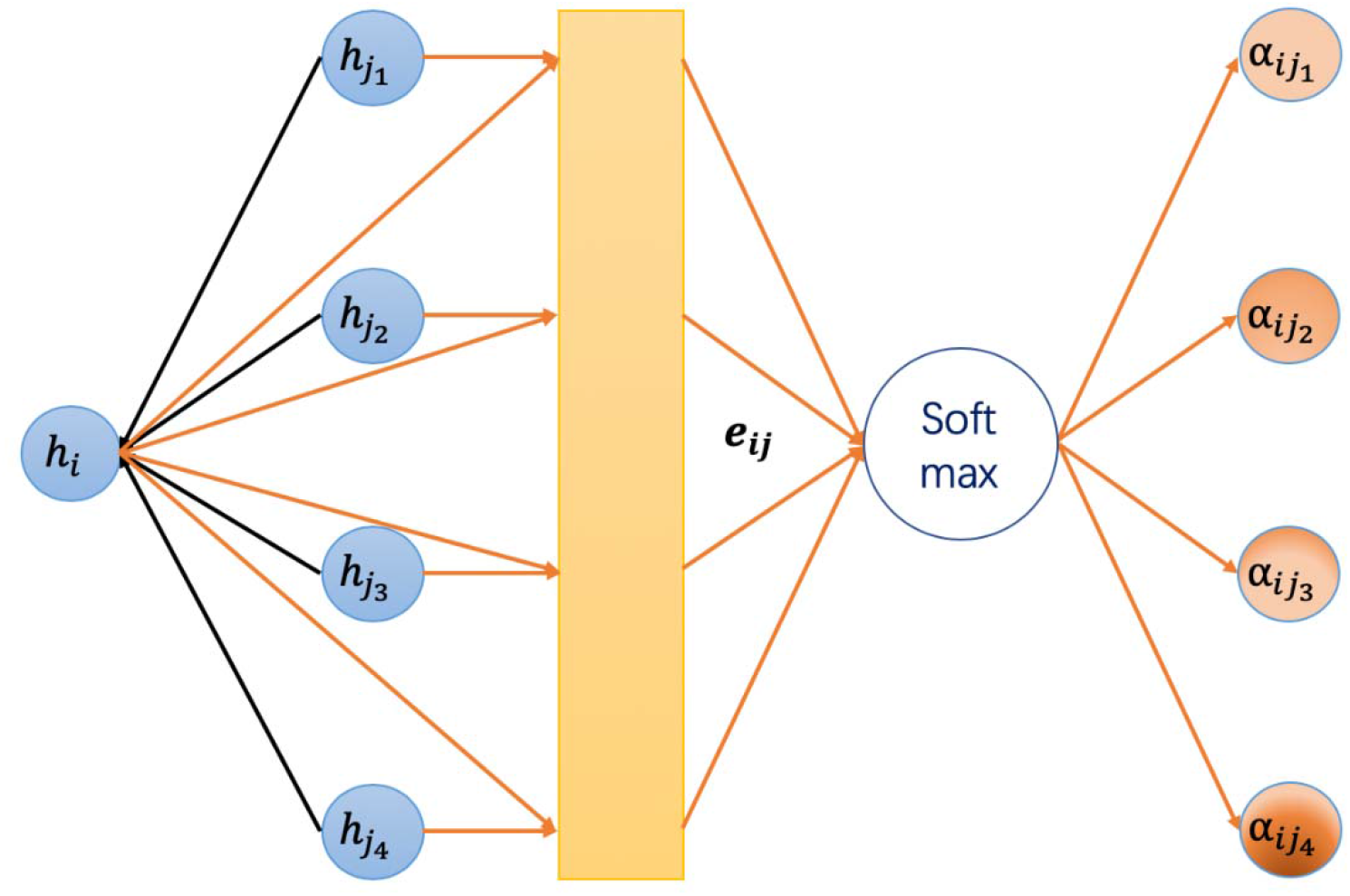

Graph Attention Networks (GATs) are a type of GNN that introduce the concept of attention mechanism to learn the importance of neighboring nodes while aggregating node representations. GATs use attention mechanisms to compute weights for the neighboring nodes, which allows the model to focus on the most relevant nodes during message passing. GATs have several advantages over traditional GNNs such as Graph Convolutional Networks (GCNs), including the ability to handle graphs with varying sizes and structures, capture both local and global graph structure information, and achieve state-of-the-art performance on various graph-related tasks.

##### 2.2.2.4 Graph Transformers

Graph Transformers are a recent development in the field of graph neural networks that extend the concept of transformer models to graph data^13^. Graph Transformers use self-attention mechanisms to compute node embeddings by attending to neighboring nodes and learn contextual representations of the graph structure. Graph Transformers have shown promising results in various graph-related tasks such as node classification, graph classification, and graph generation. They have the advantage of being able to handle graphs with varying sizes and structures, capture both local and global graph structure information, and learn hierarchical representations of the graph.

#### 2.1.5 Model Evaluation

To validate the performance of the model developed in this study, the data was split into three sets using the following format: 70% for training, 20% for validation, and 10% as a holdout test set. The accuracy, sensitivity, and specificity were calculated for both the training and test sets to assess the robustness of the model. To optimize the model performance, Weights and Biases (wandb) was used for tracking and hyperparameter tuning. wandb is a tool that helps to manage machine learning experiments, enabling the tracking of model performance metrics, as well as the hyperparameters that were tuned during training. This approach ensures that the model performance is optimized and validated in a rigorous and transparent manner.

## 4. RESULTS AND DISCUSSION

### 3.1 Visualization of Gene Expression Variation

To analyze dissimilarities between individual mRNA-Seq libraries based on gene variation, we applied Multidimensional scaling (MDS) to the integrated ML dataset. The resulting two-dimensional plot displays each dataset as a point on the x- and y-axes, with the Euclidean distance between them transformed. We focused on the top 100 genes based on log2-normalized standard deviation, and our analysis revealed that there is an overall similarity in gene expression between NAFLD and NASH, as shown in Figure 6.

**Figure 6.**
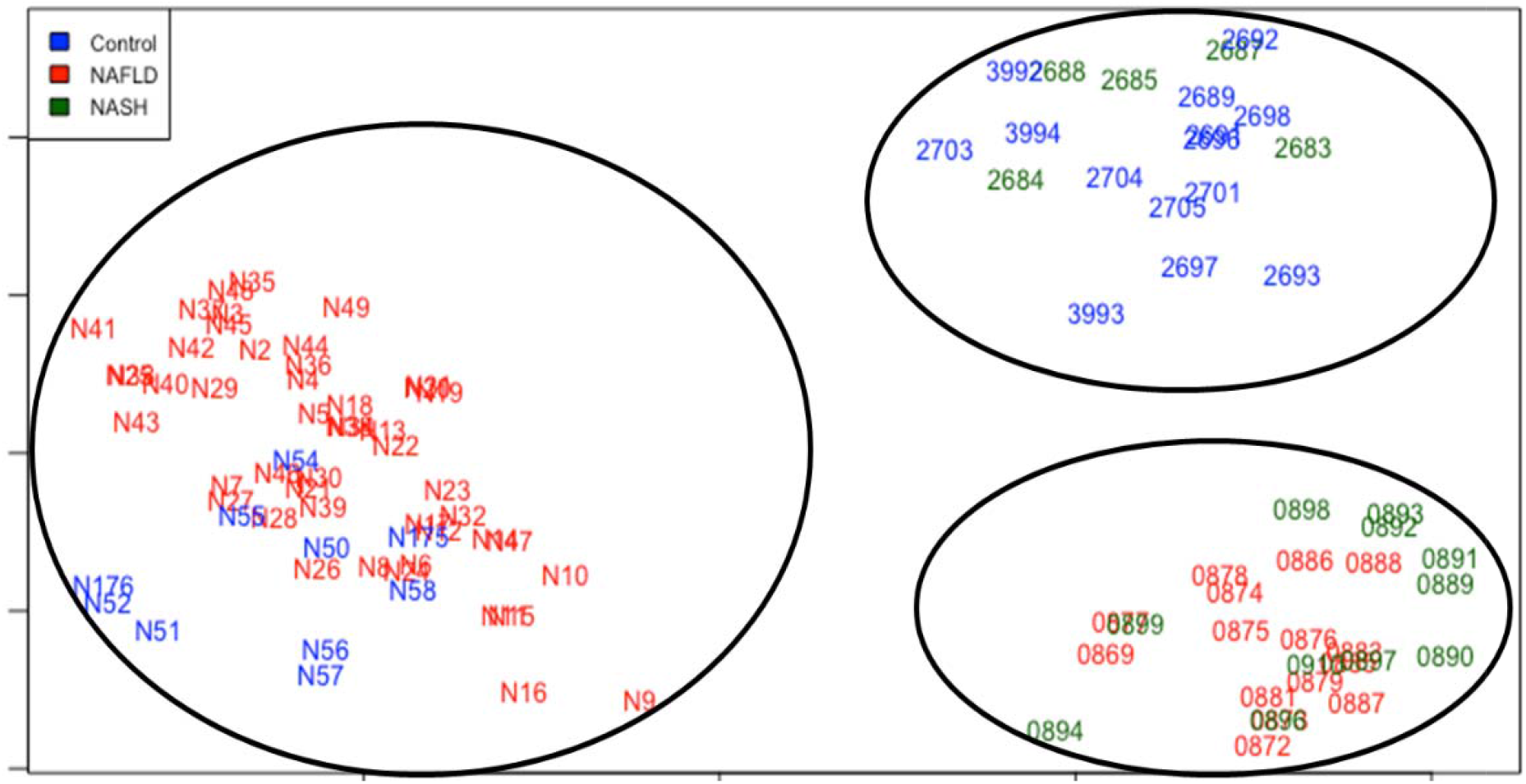
MDS plot of 300 datasets utilized for ML classification. Clustering was performed with Euclidian distances across the top 100 genes based on log2 standard deviation. Datasets are demarked by color, representing NAFLD/NASH and healthy controls.

Although there is a minor overlap between Control and NASH, as well as Control and NAFLD, this can be explained by the progressive scale of NAFLD. Despite these differences, our findings suggest that there are significant similarities in gene expression patterns between NAFLD and NASH, indicating potential common underlying mechanisms in the two conditions.

Each dataset underwent gene classification using the top ML sparse model in each etiologic-specific test group, and a heat map was generated for each dataset (Fig. 7) to visualize the results. Unsupervised hierarchical clustering was applied to the expression patterns within each column, revealing similarities in the degree of progression of NAFLD. The gene expression data showed distinct clustering between the three groups (NAFLD, NASH, and healthy). Interestingly, the clustering of genes associated with both NAFLD and NASH was observed. The control columns, however, demonstrated a clear absence of the expression of the top 100 genes. Overall, these findings suggest that the identified gene classifiers may have diagnostic or therapeutic potential for NAFLD and NASH.

**Figure 7:**
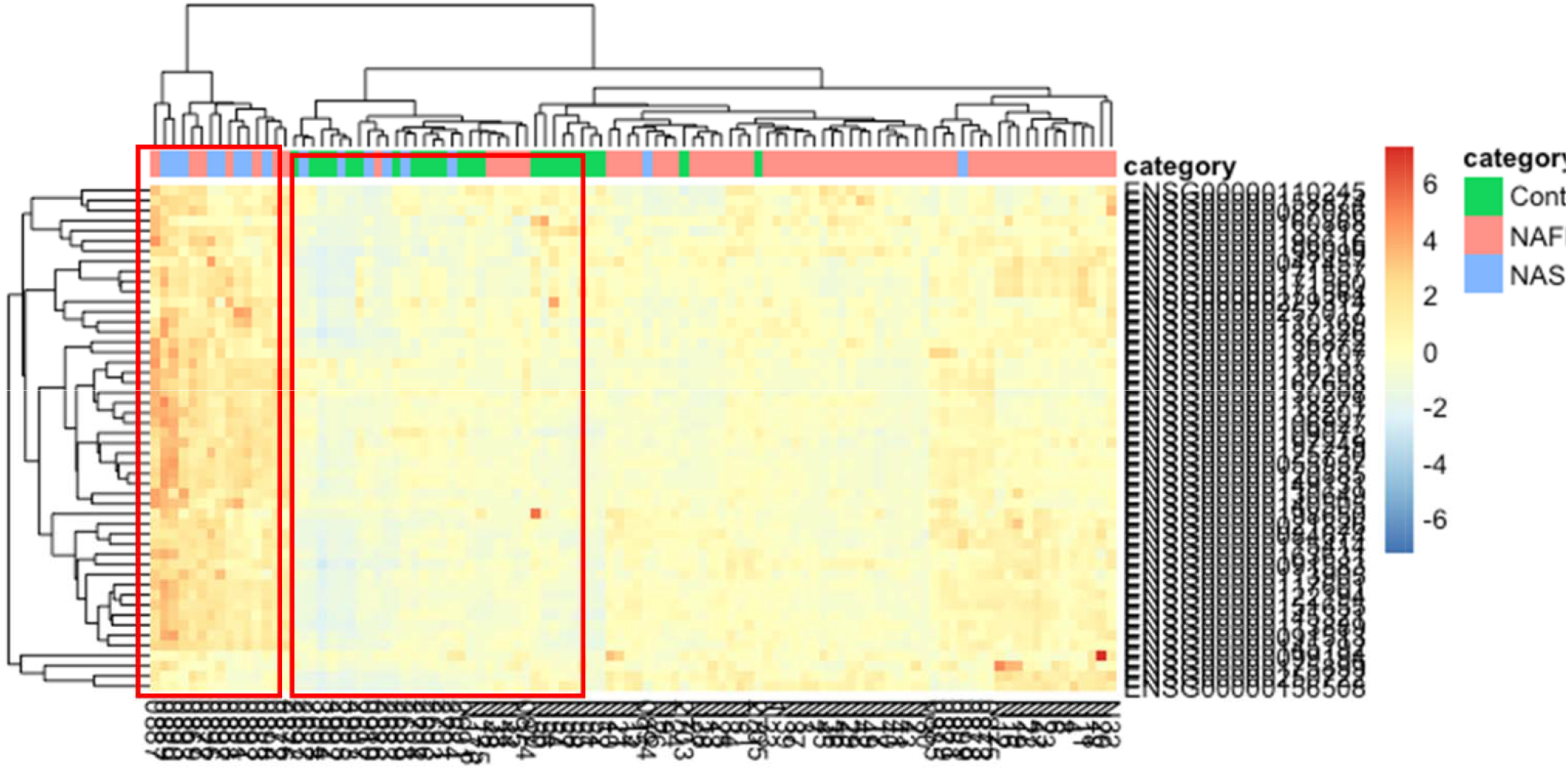
Heatmap of the 100 unique genes identified by top ML sparse classifier. Ward clustering of datasets and gene expression was performed with Euclidian distance and Pearson correlation coefficients, respectively. Visual expression modules (1, 2 or 3) were empirically identified by class dissimilarity. Clustering of samples (datasets) is more apparent for control versus disease status.

### 3.2 Baseline Machine Learning Models

Several ML models were evaluated, including Poisson linear discriminant analysis (PLDA), sparse Poisson linear discriminant analysis with a power transformation (PLDA2), negative binomial linear discriminant analysis (NBLDA), support vector machine (SVM); nearest shrunken centroids (NSC), voom-based nearest shrunken centroids (voom-NSC). The results of the machine learning (ML) performance evaluation across all testing sets are shown in Table 2. Among these models, SVM demonstrated the highest performance in terms of balanced accuracy, sensitivity, and specificity.

**Table 2.** ML performance across all testing sets. Calculations for ML performance metrics are defined by Goksuluk and colleagues (2019).

| Baseline Models | Accuracy |  | Sensitivity |  | Specificity |  |
| --- | --- | --- | --- | --- | --- | --- |
|  | Training (%) | Testing (%) | Trainin g (%) | Testing (%) | Trainin g (%) | Testing (%) |
| SVM | 74.53 | 76.92 | 86.21 | 90.00 | 98.36 | 58.06 |
| NSC | 66.98 | 62.64 | 75.86 | 63.33 | 95.63 | 67.74 |
| voomNSC | 68.87 | 58.24 | 86.21 | 56.67 | 92.9 | 67.74 |
| PLDA | 62.09 | 58.24 | 67.86 | 61.67 | 93.44 | 67.74 |
| PLDA2 | 68.54 | 61.54 | 76.67 | 63.33 | 96.17 | 67.74 |
| NBLDA | 71.6 | 63.74 | 73.68 | 65.00 | 93.55 | 67.74 |

It is worth noting that SVM is a non-sparse classifier. This means that it does not employ built-in algorithms for feature selection, unlike other sparse classifiers. Instead, SVM uses the kernel trick to transform data into a higher-dimensional space where it can be more easily separated by a hyperplane. This makes SVM a powerful and flexible model for solving complex classification problems with large datasets. On the other hand, sparse classifiers rely on selecting a subset of the available features, either through regularization or feature selection algorithms. This approach can be useful in situations where the number of features is large and the signal-to-noise ratio is low, but it can also result in oversimplified models that may underperform on unseen data.

In Figure 8, Venn diagram displays the number of genes identified by each ML model and the degree of overlap between them. This can provide insight into which models are most effective at identifying similar sets of genes and can aid in the selection of the best model for a given dataset. In particular, the high degree of overlap among the genes identified by NBLDA, PLDA2, and voom NSC suggests that these models may be more robust and consistent in their results compared to other models. Table 3 provides additional details on the top 17 genes identified using sparse classifiers, which typically use feature selection methods to identify a subset of the most informative genes for prediction. These genes were considered as potential biomarkers or targets for further investigation. In-lab validation using qPCR studies were performed.

**Figure 8:**
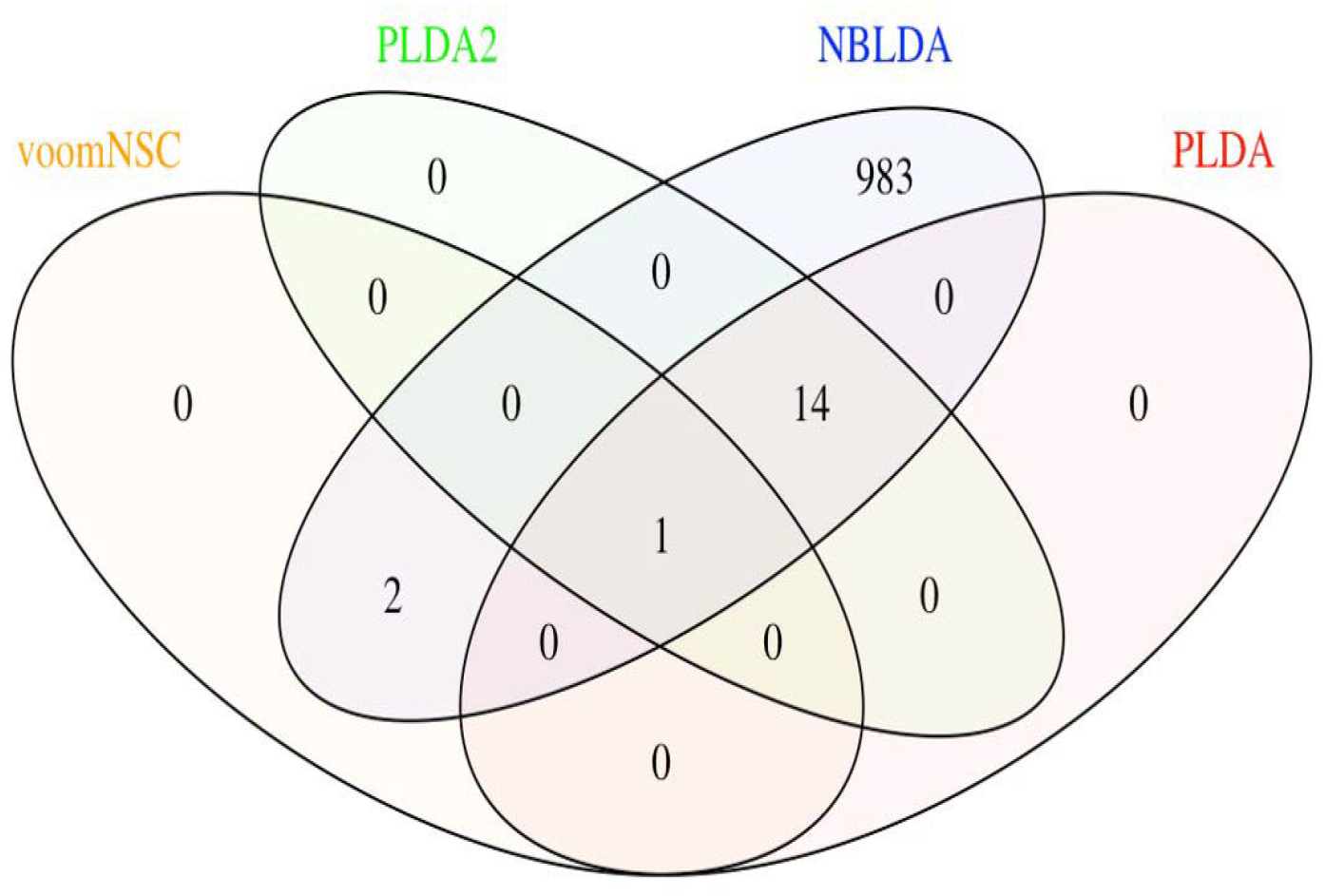
Venn diagram illustrating the overlap of the genes identified using various ML models.

**Table 3:** Top 17 genes identified using baseline ML models.

| S No | Ensembl ID | Gene |
| --- | --- | --- |
| 1 | ENSG00000198840 | MT-ND3 |
| 2 | ENSG00000109072 | VTN |
| 3 | ENSG00000118137 | APOA1 |
| 4 | ENSG00000130208 | APOC1 |
| 5 | ENSG00000140505 | CYP1A2 |
| 6 | ENSG00000171759 | PAH |
| 7 | ENSG00000234745 | HLA-B |
| 8 | ENSG00000119711 | ALDH6A1 |
| 9 | ENSG00000183044 | ABAT |
| 10 | ENSG00000182492 | BGN |
| 11 | ENSG00000163682 | RPL9 |
| 12 | ENSG00000171863 | RPS7 |
| 13 | ENSG00000125844 | RRBP1 |
| 14 | ENSG00000100342 | APOL1 |
| 15 | ENSG00000164587 | RPS14 |
| 16 | ENSG00000092841 | MYL6 |
| 17 | ENSG00000166278 | C2 |

### 3.3 XAI methods

Different attention-based models for feature selection in gene expression data to identify potential RNA biomarkers associated with nonalcoholic fatty liver disease (NAFLD) and nonalcoholic steatohepatitis (NASH) were evaluated. Evaluated the performance of four different attention-based models, including a Single Head Attention Model, a Multi-Head Attention Model, a Graph Attention Network (GAT) Model, and a Graph Transformer Model.

To evaluate the performance of these models, we trained them on our preprocessed count matrix data and tested them using a hold-out test set. The training accuracy and testing accuracy for each model and recorded the results in the table below (Table 4).

**Table 4.**
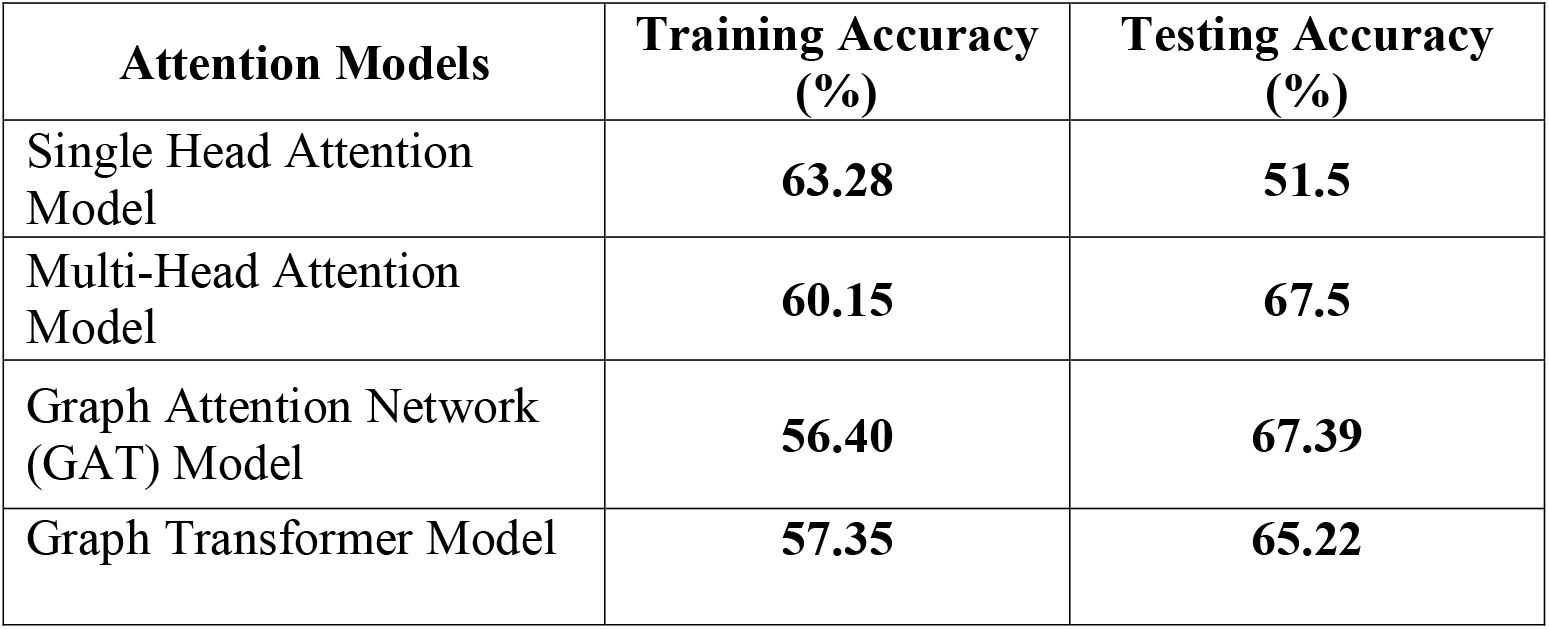
Accuracy of attention models.

The results showed that the Multi-Head Attention Model achieved the highest testing accuracy, with a score of 67.5%. The Single Head Attention Model had the lowest testing accuracy, with a score of 51.5%. The Graph Attention Network and Graph Transformer Models both achieved moderate testing accuracy, with scores of 67.39% and 65.22%, respectively.

Overall, our evaluation of these attention-based models provided insight into their potential for identifying RNA biomarkers associated with NAFLD and NASH. The Multi-Head Attention Model showed the most promising results, and we planned to explore it further and compare its performance to the baseline variance filtering method in our future experiments.

Figure 9 shows, the top 20 RNA biomarkers identified using attention mechanism. Genes with highest attention scores in our analysis were found to be RBM6, GCKR, and ADIPOR2. These biomarkers have been previously implicated in various biological processes, and their clinical significance has been supported by a growing body of literature. These biomarkers have been previously linked to a range of biological processes, such as RNA splicing, glucose metabolism, and adipocyte function, respectively. Moreover, a number of studies have reported their potential clinical significance in various diseases, including cancer, diabetes, and obesity.

**Figure 9:**
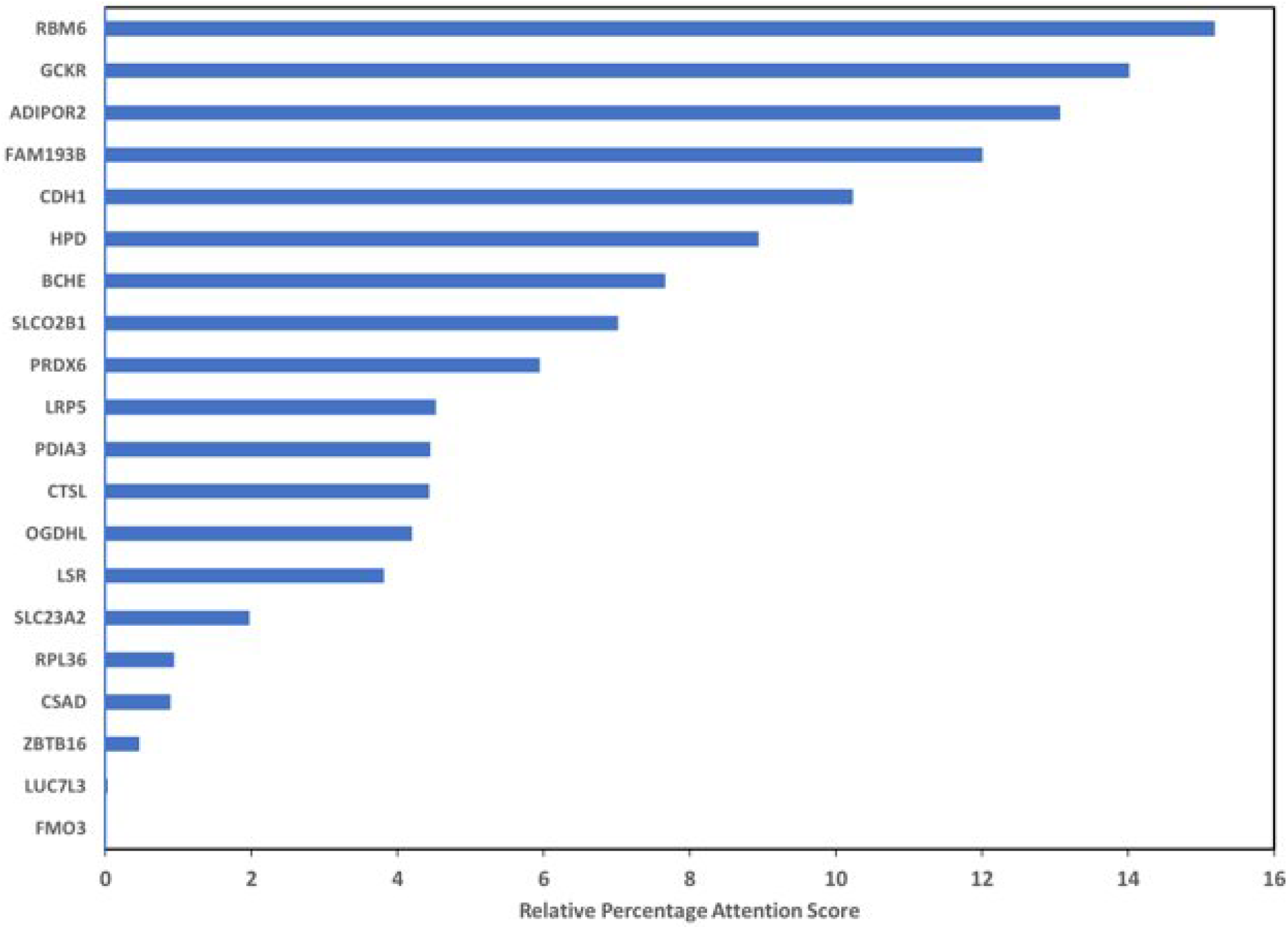
Attention scores of top 20 biomarkers.

Overall, the identification of these biomarkers with high attention scores provides valuable insights into the molecular mechanisms underlying disease development and progression and may ultimately lead to the development of more accurate diagnostic and therapeutic strategies.

Overall, our analysis of gene expression data provided insight into potential RNA biomarkers associated with NAFLD. The higher relative expression levels of HLA-B, APOL-1, and MT-ND3 in individuals with NAFLD suggest that these genes may play a role in the development and progression of the disease. Further validation and investigation of these potential RNA biomarkers will be necessary to determine their usefulness in clinical applications.

## 5. CONCLUSIONS

In conclusion, the study found that machine learning approaches can be effective in identifying genes that are differentially expressed in patients with NAFLD and NASH. Baseline machine learning approaches identified 17 genes that showed differential expression in these patients. Furthermore, preliminary data from in-lab validation suggest upregulation of MT-ND3, APOC1, HLA-B, APOL-1, MT-ND3 in these patients.

The study also found that attention-based approaches, specifically attention/XAI, showed promise in improving the prediction of specific biomarkers. Both baseline and attention-based approaches were able to differentiate genomic data of patients with NAFLD/NASH vs. control. These findings suggest that attention-based approaches may provide a more accurate and precise analysis of genomic data.

Overall, this study highlights the potential of machine learning approaches to identify biomarkers for the diagnosis and treatment of NAFLD and NASH.

